# Sensorimotor training lightens the perceived weight of body augmentation devices

**DOI:** 10.64898/2026.04.17.718984

**Authors:** Dominika Radziun, Arina Schippers, Matthew R. Longo, Luke E. Miller

## Abstract

Wearable haptic and robotic devices must transmit useful forces and tactile information without remaining perceptually intrusive. Here we show that perceived weight provides a behavioral marker of this transition toward transparency. Participants judged the weight of their hand with or without a finger-extending exoskeleton before and after object-manipulation training with either the exoskeleton or a structurally matched control device. Psychometric modelling decomposed perceived effector weight into biological-hand and artificial-device components. Attachment produced baseline attenuation of both hand and device weight, whereas training selectively increased attenuation of the finger-extending exoskeleton. Thus, active use made the augmenting device contribute less to perceived effector weight while leaving perceived hand weight stable. These findings suggest that wearable devices become perceptually lighter when their mechanical consequences are integrated into sensorimotor control. Perceived weight offers a compact metric for evaluating transparency, embodiment, and usability in wearable haptic and robotic interfaces.

**Summary:** Sensorimotor training makes wearable finger-extending exoskeletons feel lighter.

## INTRODUCTION

Wearable haptic and robotic interfaces are increasingly designed to operate as physically coupled extensions of the body. Fingertip tactors, force-feedback gloves, exoskeletons, prosthetic interfaces, and supernumerary robotic limbs can provide tactile or force information during virtual interaction, teleoperation, rehabilitation, and human augmentation (*1–5*). Their success depends not only on the fidelity of the information they deliver, but also on whether the device can recede from awareness during action, remaining transparent in the sense of not becoming perceptually intrusive or attention-demanding. A wearable interface that is mechanically effective but perceptually salient may remain uncomfortable, distracting, or effortful to use. Thus, a central challenge for haptics and wearable robotics is to design devices that are informative when needed but otherwise unobtrusive to the user, a goal long emphasized in haptic interface and teleoperation research (*6*).

A distinctive feature of ordinary bodily experience is this form of transparency. During skilled action, our limbs typically recede from awareness and function as the medium of interaction rather than as perceptual objects (*7*). This transparency is reflected in systematic perceptual biases: humans reliably underestimate the weight of their own hands relative to their actual physical mass (*8*). Such underestimation may reflect predictive motor processes that attenuate the sensory consequences of the body’s own mechanical load, consistent with broader evidence that self-generated tactile sensations are attenuated during action (*9, 10*). In this view, the hand feels lighter than its physical weight because its forces are expected, controlled, and integrated into the internal models used for action.

Wearable robotic devices challenge the limits of this perceptual transparency. Unlike handheld tools, wearable devices are attached directly to the body and contribute forces through the same somatosensory channels as the biological limb. Exoskeletons and supernumerary robotic limbs also change the functional morphology of the acting body, requiring users to learn new mappings between movement, contact, and task outcome (*3, 11–13*). These devices therefore pose a problem that is both mechanical and perceptual: their added mass must be controlled, but it must also become sufficiently integrated into body representation that it does not remain salient as an external load (*14, 15*).

Evidence from prosthetics suggests that perceived weight is sensitive to embodiment. Prosthetic limbs can feel heavier than biological limbs even when they are physically lighter, and restoring sensory feedback can reduce perceived prosthesis weight by promoting embodiment (*16*). Prostheses, however, substitute for an absent or impaired biological effector. Wearable augmentation devices pose a different challenge because they are added to an intact body. The nervous system must therefore represent the biological limb and the artificial structure together, while preserving their distinct mechanical contributions (*17*). Whether the perceived weight of such an augmenting device can be selectively attenuated through sensorimotor experience remains unknown.

This question is important for haptics and robotics because perceived heaviness may influence comfort, usability, and long-term adoption independently of physical mass. Haptic devices are commonly evaluated in terms of task performance, feedback fidelity, subjective comfort, or embodiment (*2, 18–20*). These measures are essential, but they do not directly quantify whether the device’s own mechanical presence has become transparent. Perceived weight offers a behavioral and theoretically meaningful measure of this process. If the forces generated by an attached device are integrated into sensorimotor predictions, the device should contribute less to perceived effector weight even though its physical mass is unchanged.

Here, we tested whether sensorimotor training with wearable finger-extending exoskeletons (*11*) changes their perceived weight. Finger-extending exoskeletons offer a tractable model for studying wearable body augmentation, allowing us to test how artificial structures attached to the body become integrated into sensorimotor control in ways that may generalize to broader haptic and robotic technologies. Participants judged the weight of their right hand, either alone or while wearing a finger-extension device, before and after object-manipulation training. A structurally matched non-augmenting control device allowed us to separate effects of general device exposure from effects specific to functional augmentation.

Because wearable devices are perceived as part of a combined biological-artificial effector, changes in perceived hand-plus-device weight could reflect altered perception of the hand, the device, or the combined effector. To resolve this ambiguity, we developed a component model in which perceived effector weight is expressed as the sum of biological-hand and artificial-device contributions, each scaled by an attenuation parameter. This allowed us to test whether sensorimotor training selectively reduced the perceived contribution of the wearable device itself.

We hypothesized that active sensorimotor training would selectively increase attenuation for the finger-extending exoskeleton because it changes finger morphology, object-contact geometry, and the mapping between hand movement and task outcome. Under this account, attenuation of the artificial component provides a behavioral marker of wearable haptic transparency: as the device becomes integrated into sensorimotor control, its physical mass contributes less to perceived effector weight.

## RESULTS

### Experimental design and training procedure

Participants performed a weight-comparison task (*8*) to estimate the perceived weight of the right-hand effector, either as the biological hand alone or as the hand plus a wearable body-augmentation device (Fig. 1A). The finger-extension device was designed to alter functional hand morphology by extending each finger while preserving active control of the biological fingers; its numbered structural components are shown in Fig. 1B. Measurements were collected before and after sensorimotor training with either the finger-extending exoskeleton or a structurally matched non-augmenting control device (Fig. 1A). On each trial, participants compared the perceived weight of the right-hand effector with a reference weight suspended from the opposite wrist. This design allowed us to measure whether short-term training changed perceived effector weight, and whether any change depended on training with a functionally augmenting device. Participants completed object-manipulation training while wearing their assigned device (Fig. 1D). Training involved grasping and manipulation tasks designed to expose participants to the device’s mechanical consequences during active use.

**Figure 1.**
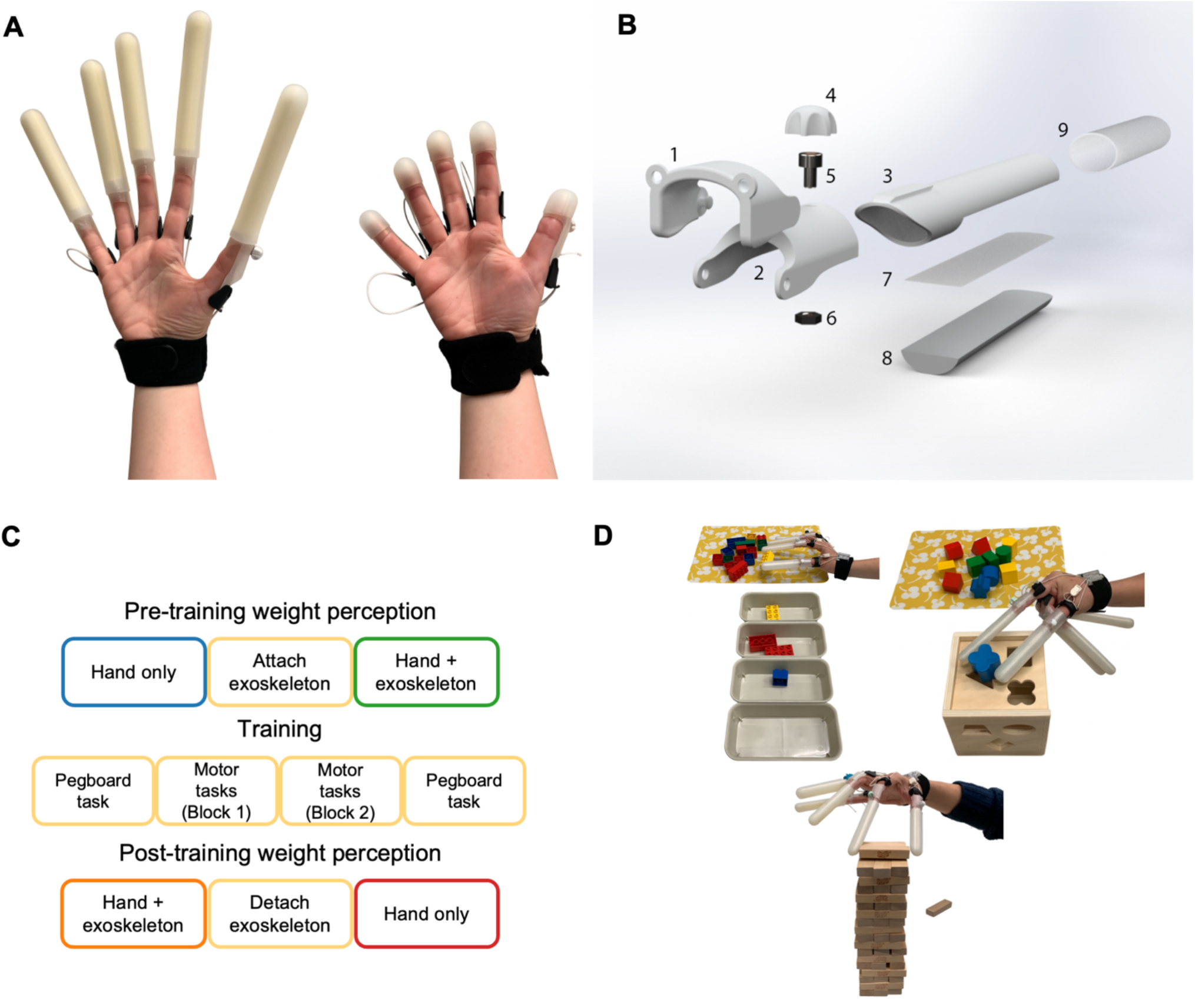
Device structure and experimental design. (A) The finger-extending exoskeleton elongated each finger by approximately 10 cm while preserving native finger-joint mobility. A structurally matched control device had the same general attachment format but did not meaningfully extend the fingers. Participants trained with one of the two devices. (B) Structural components of the finger-extension device. Numbered labels indicate: 1, proximal segment with joint and elastic-band connection; 2, intermediate segment with joint and adjustable length to the fingertip; 3, fingertip cap; 4, knob for securing the fingertip cap; 5, fastening screw for fingertip attachment; 6, embedded nut for fingertip fastening; 7, double-sided tape securing the internal padding; 8, foam padding providing a comfortable contact surface with the biological fingertip; and 9, silicone sleeve forming the external gripping surface. (C) Experimental design. Participants judged perceived hand weight under two effector conditions, hand only and hand plus device, before and after sensorimotor training. (D) During training, participants performed object-manipulation tasks while wearing either the finger-extending exoskeleton or the control device; performance was quantified by task completion time or number of completed actions.

### Theoretical model of weight attenuation

To gain traction on partitioning participant behavior into hand and exoskeleton-specific effects, we developed a theoretical model of weight attenuation. The weight of a person’s hand *w*_*h*_ exerts forces on their upper limb that are picked up by sensory receptors (e.g., mechano- and proprioceptors) and communicated to their nervous system as sensory feedback (*21–23*). Previous research demonstrates that the weight of the hand can be perceived, though it is underestimated. We can formalize this with the following linear equation:

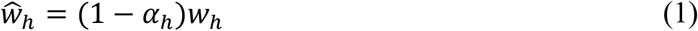

Where 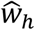 is the perceived weight of the hand and *α*_*h*_ is the attenuation factor. When the attenuation factor is zero, perception is veridical; perception is underestimated when *α*_*h*_ is greater than zero and overestimated when *α*_*h*_ is less than zero.

When a wearable device is attached to the hand—as in the exoskeleton of the present experiment—the forces picked up by the sensory receptors now include the weight of the device: *w*_*he*_ = *w*_*h*_ + *w*_*e*_. If we assume that this linearly alters the perceived weight, then we augment Equation 1 as follows:

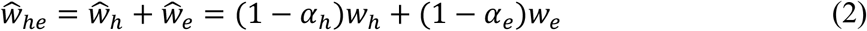

Where 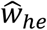 is the total perceived weight of the hand and exoskeleton combined, 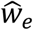 is the perceived weight of the exoskeleton alone, and *α*_*e*_ is the exoskeleton-specific attenuation factor. Note that this formalization of 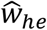 assumes that the nervous system can make unique estimates of hand and exoskeleton weight, perhaps via distinct internal models of the unique forces exerted by the hand and exoskeleton on the upper arm (*11, 24*).

### Model-based decomposition of perceived weight

The present experimental paradigm measured 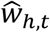 and 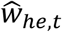 at two distinct time points *t*: before (*t* = 1) training (i.e., the baselines); and after (*t* = 2) sensorimotor training with either the exoskeleton or control device (n=17 per device). We simultaneously fit psychometric functions to trial-wise judgments in all four conditions in order to estimate the attenuation parameters from Equations 1 and 2. We then used a model-based decomposition approach to isolate distinct components of these attenuation parameters and their dynamics.

We hypothesized that the perceived weight of the hand and exoskeletons across time can be explained by three latent variables. Before training, perceived weight (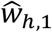 and 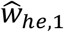) is only affected by the baseline attenuation parameters, *α*_*h*,1_ and *α*_*e*,1_. The effect of training on the attenuation parameters is further formalized as follows, with two additional terms:

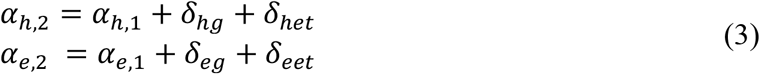

Where *δ*_*hg*_ and *δ*_*eg*_ reflect a general change to hand and exoskeleton attenuation, respectively; the value of this parameter is independent of the device used during training and may be driven by practice or persistent movement. In contrast, *δ*_*het*_ and *δ*_*eet*_ reflect a change to perceptual attenuation that is driven specifically by training with the exoskeletal device. That is, when training with the exoskeletal device, the value of this parameter can be non-zero; when training with the control device, its value is fixed at zero and can therefore be dropped from Equation 3. Thus, the model tested whether active use of the finger-extending device selectively reduced its contribution to perceived effector weight, while leaving perceived biological hand weight comparatively stable. Given the subject-level fits of our psychometric functions, we used linear-mixed modelling to isolate all variables from Equations 1-3. See the Methods section for more mathematical details of the modelling.

### Attachment attenuates perceived hand and device weight

We first analyzed the weight attenuation parameters for the perceived weight of the hand (*α*_*h*,1_) and exoskeletal device (*α*_*e*,1_). We found substantial perceptual attenuation in both conditions (Fig. 2A), indicating that perceived weight was reduced relative to physical weight. Consistent with prior findings (*8, 25*), we observed significant attenuation for the perceived weight of the hand (27.3%, Bayesian bootstrap SD = 6.0%; LMM: t(65) = 4.41, p < 0.001; BF_10_ = 87.48–399.89 across prior widths). Furthermore, we also observed significant attenuation for the perceived weight of the exoskeleton (25.0%, Bayesian bootstrap SD = 9.8%; LMM: t(65) = 2.45, p = 0.017; BF_10_ = 3.51– 16.07). Interestingly, the magnitude of attenuation was similar for both hand and exoskeleton. The estimated difference between baseline hand and exoskeleton attenuation was small (2.3%, Bayesian bootstrap SD = 11.5%; LMM: t(65) = -0.19, p = 0.846; BF_10_ = 0.22–1.00), providing little evidence that baseline attenuation differed between the biological and artificial components. This suggests that attachment of the device was sufficient to alter perceived effector weight at an early stage.

**Figure 2.**
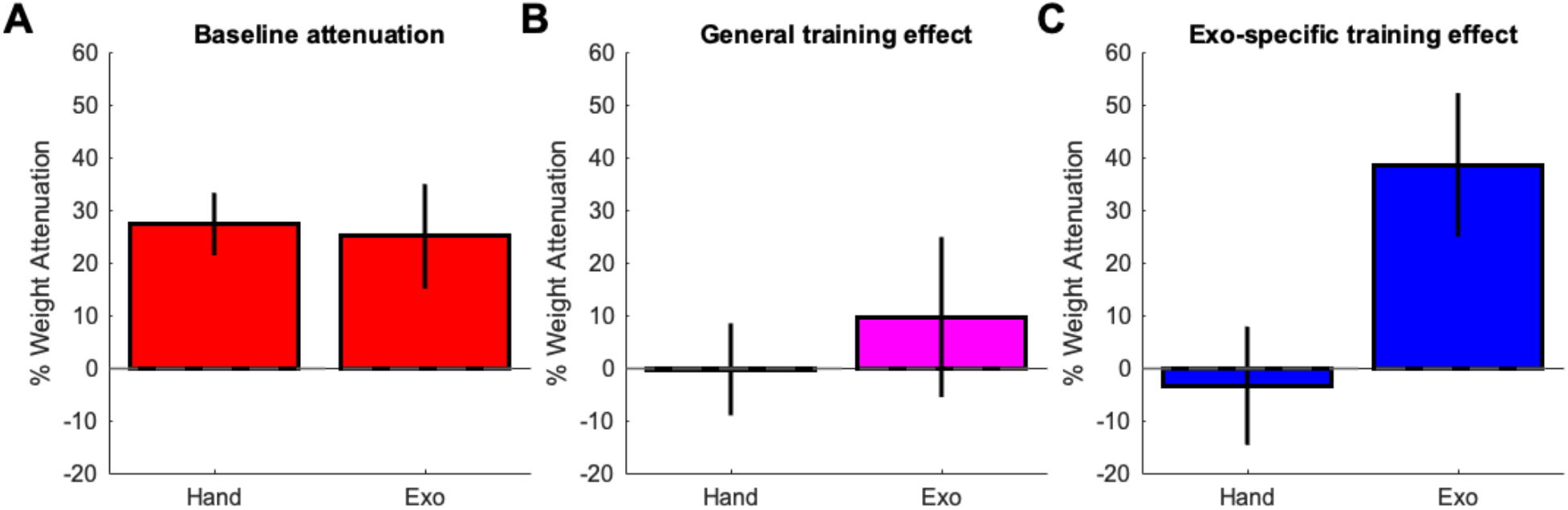
Model-based decomposition of perceived weight attenuation. (A) Baseline attenuation parameters represent the contribution of hand and device weight to perceived weight before training. (B) General training-effect parameters capture changes in perceived hand and device weight after training across both groups, independent of the device used during training. (C) Finger-extension-specific parameters represent additional post-training changes in the finger-extension group relative to the control-device group. Bars show the magnitude of the best-fitting coefficients, expressed as percentage attenuation, describing how physical hand and device mass contributed to perceived weight judgments before and after training. Error bars denote the standard deviation of the Bayesian bootstrap distribution estimated from 5,000 resamples. Statistical tests reported in the text were obtained from linear mixed-effects models fit to the same participant-level attenuation parameters. The parameter pattern indicates baseline underestimation of both hand and device weight, little evidence for a general training-related change in attenuation, and an additional lightening effect specific to training with the finger-extending exoskeleton.

### Training selectively lightens the augmenting device

We next investigated the effect of training on attenuation (Equation 3), where changes can either be general to the use of both devices (δ_g_) or specific to the exoskeleton training (δ_et_). Whether training modulated attenuation depended on the functional properties of the device. We found no evidence for a general effect of training on the perceived weight of the hand (−0.3%, Bayesian bootstrap SD = 6.0%; LMM: t(65) = 0.27, p = 0.790; BF_10_ = 0.11–0.48) or the exoskeleton (9.7%, Bayesian bootstrap SD = 12.7%; LMM: t(65) = 0.94, p = 0.350; BF_10_ = 0.32–1.47; Fig. 2B), indicating that mere exposure or repeated use was not sufficient to reliably modulate attenuation. In contrast, training with finger-extending exoskeletons strongly modulated their perceived weight (38.6%, Bayesian bootstrap SD = 12.4%; LMM: t(65) = 2.62, p = 0.011; BF_10_ = 23.59–107.83; Fig. 2C), indicating that attenuation increased specifically when active use involved an augmenting device that altered finger morphology and action kinematics rather than active use of a wearable device per se. After sensorimotor training, the physical mass of the finger-extending exoskeleton contributed less to perceived effector weight than it did before training: the device was perceived as lighter relative to its actual mass. This effect was specific to the exoskeleton, as the corresponding exoskeleton-specific parameter for the biological hand was close to zero (−3.4%, Bayesian bootstrap SD = 8.3%; LMM: t(65) = -0.90, p = 0.371; BF_10_ = 0.16–0.71), and the device-specific effect was larger for the exoskeleton than for the biological hand (LMM difference: 41.1%, SE = 16.8%; t(65) = 2.44, p = 0.017; BF_10_ = 29.69–135.74).

### Condition-level estimates support the model-based result

We also performed a complementary condition-level analysis directly on the psychometric fits, which yielded a consistent pattern of results. For this descriptive condition-level analysis, negative values indicate underestimation relative to physical weight. Participants underestimated biological hand weight both before and after training in both groups, with no reliable pre-post change in either the finger-extension group (pre: −27.7% ± 9.2 SEM; post: −21.5% ± 9.0 SEM; t(16) = 0.95, p = 0.358) or the control-device group (pre: −19.7% ± 7.4 SEM; post: −27.2% ± 6.8 SEM; t(16) = - 1.12, p = 0.280). In the hand-plus-device condition, attenuation increased after training in the finger-extension group (pre: −30.6% ± 7.3 SEM; post: −40.6% ± 6.7 SEM; t(16) = -2.35, p = 0.032), with a weaker threshold-level change in the control-device group (pre: −22.7% ± 6.6 SEM; post: −29.4% ± 6.8 SEM; t(16) = -2.12, p = 0.050). When the device-related contribution was inferred by subtracting the hand-only estimate from the hand-plus-device estimate, attenuation increased substantially after training in the finger-extension group (pre: −36.4% ± 14.9 SEM; post: −82.4% ± 9.5 SEM; t(16) = -2.93, p = 0.010), but not in the control-device group (pre: −30.2% ± 14.1 SEM; post: −35.6% ± 12.1 SEM; t(16) = -0.34, p = 0.741). This analysis supported the regression-based conclusion that active training selectively reduced the perceived contribution of the augmenting exoskeleton to total effector weight.

### Practice improves control of the finger-extending exoskeleton

The initial performance cost of using the finger-extending exoskeleton diminished substantially with practice. As shown in Fig. 3, participants wearing the exoskeleton were initially much slower on the pegboard-transfer tasks, but their completion times decreased markedly during the second training block. This improvement was evident for both large-peg transfer (Wilcoxon V = 0.0, p < 0.001, BF_10_ = 3970.03) and small-peg transfer (V = 11.0, p = 0.001, BF_10_ = 38.64). Participants wearing the control device also improved, although their performance began closer to its later level and the change across blocks was less pronounced, particularly for the small pegs (large pegs: V = 0.0, p < 0.001, BF_10_ = 1502.34; small pegs: V = 28.0, p = 0.012, BF_10_ = 2.89). Thus, training produced especially pronounced gains when participants had to adapt their movements to the altered finger morphology of the augmenting device. Results for the remaining training tasks are reported in the Supplementary Materials and shown in Supplementary Fig. S1.

**Figure 3.**
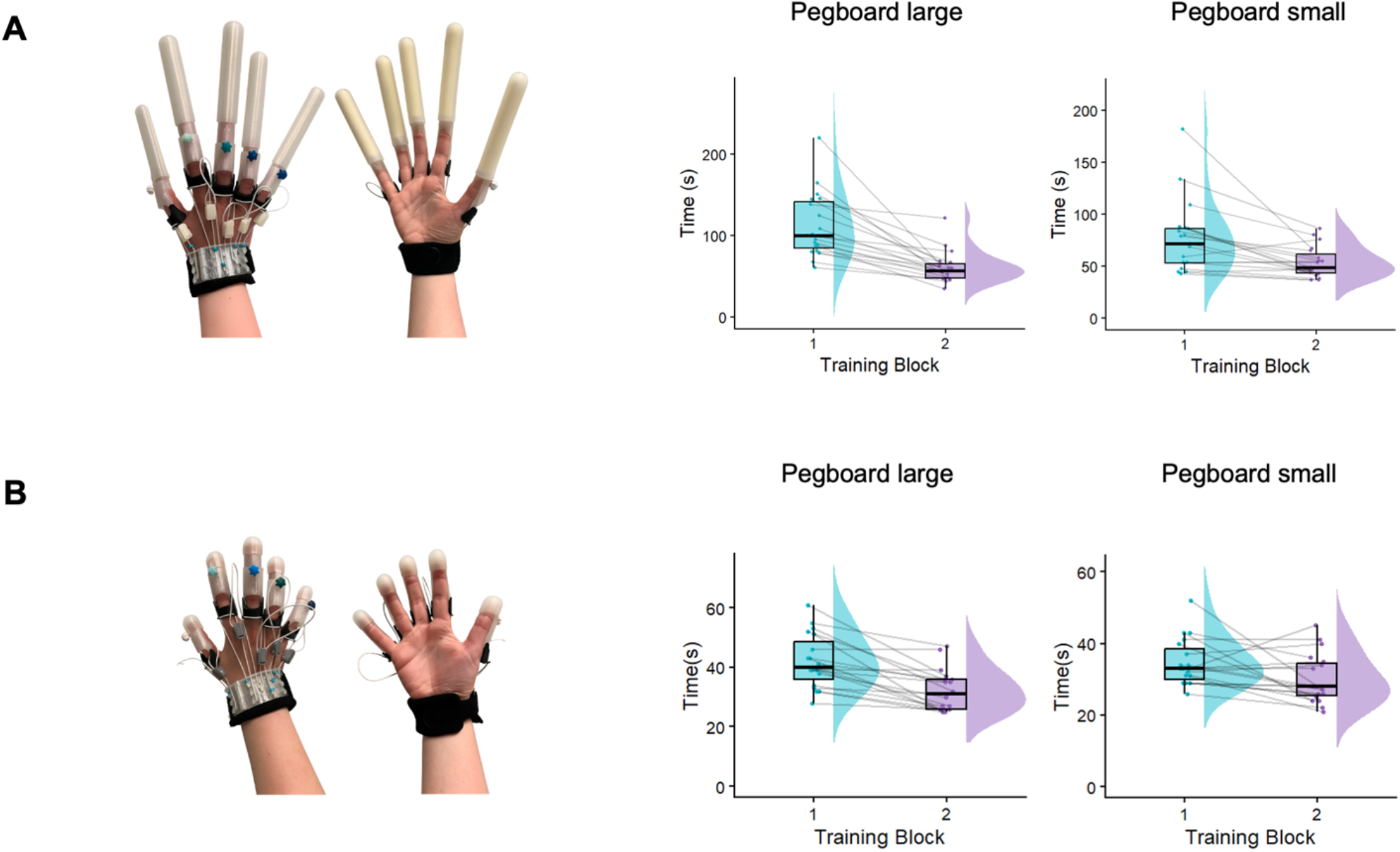
Pegboard-transfer performance following sensorimotor training. Participants completed large-peg and small-peg transfer tasks while wearing either the finger-extending exoskeleton or the structurally matched control device. Performance was quantified as task completion time across the two training blocks. The full set of training tasks, including color sorting, shape sorting, and Jenga-based tasks, is shown in Supplementary Fig. S1.

## DISCUSSION

These findings show that sensorimotor training makes a wearable augmentation feel lighter. Participants underestimated the weight of their biological hand, consistent with previous evidence that the hand’s physical mass is systematically attenuated in perception (*8, 25*). Critically, the attached device was also attenuated, and this attenuation increased selectively after training with the finger-extending exoskeleton. Thus, the device did not simply add veridical physical mass to the hand. Instead, its contribution to perceived effector weight depended on how it was coupled to action. Training therefore changed the perceptual consequences of wearing the device: after active use, the same physical mass contributed less to the experienced weight of the hand-device effector. This suggests that perceived heaviness is not fixed by device mechanics alone, but can be altered by sensorimotor experience.

The specificity of the effect is important. Training did not produce a comparable change in perceived biological hand weight, and the strongest device-specific lightening occurred for the finger-extending exoskeleton rather than the structurally matched control device. This pattern argues against a purely general explanation based on repeated testing, passive wearing, or motor activity. Instead, the effect appears linked to functional augmentation: the exoskeleton changed the effective length of the fingers and required participants to learn a new relationship between finger movements, object contact, and task outcomes. Such learning is consistent with the formation or updating of internal models that predict the sensory consequences of action (*26, 27*).

The model-based decomposition is central to this interpretation. By estimating separate attenuation parameters for hand and device contributions, the model showed that training selectively reduced the perceived contribution of the augmenting exoskeleton. This approach is consistent with the idea that internal models predict the sensory consequences of body and tool dynamics during action (*9, 10, 24, 26–28*). It also extends embodiment-based approaches to artificial limbs and devices by asking whether the artificial component itself becomes perceptually attenuated (*14–16*). Thus, the model provides a general framework for wearable haptics: perceptual transparency can be quantified as the extent to which an artificial component’s physical mass is attenuated after sensorimotor experience.

We interpret these findings as evidence for a progressive integration of wearable devices into sensorimotor representations and internal models. Passive attachment may initiate attenuation because the device is mechanically coupled to the body and contributes to the sensory signals generated during limb movement. This is consistent with work showing that artificial extensions and supernumerary effectors can alter body representations over relatively short timescales (*11, 12, 29*). Active sensorimotor training may then further refine internal models of the hand-device system when the device changes the user’s action capabilities. In the present study, the finger-extending exoskeleton required participants to adapt to a modified hand morphology and altered relationships between finger movement, object contact, and task outcome. This functional adaptation was accompanied by a selective reduction in perceived device weight, suggesting that the exoskeleton’s mechanical consequences became more effectively integrated into sensorimotor control.

The present results extend principles from prosthetics to wearable augmentation while also highlighting an important distinction. In prosthesis users, sensory feedback can reduce perceived prosthesis weight by promoting embodiment of the artificial limb (*16*). Here, however, the reduction in perceived device weight occurred without adding artificial somatosensory feedback. Instead, training with an intact hand-device system was sufficient to make the augmenting device feel lighter. This suggests that predictable coupling between action and device mechanics can promote sensorimotor transparency even when no new feedback channel is introduced. This distinction matters because augmentation does not replace the biological limb: the nervous system must represent biological and artificial components together. The model-based decomposition used here suggests that attenuation can be selectively assigned to the artificial component, rather than reflecting only a global change in perceived limb weight.

This selective attenuation has direct relevance for haptics and wearable robotics. Wearable haptic devices are often designed to stimulate the skin or apply forces through continuous contact with the body (*2, 4*). In teleoperation and virtual interaction, haptic transparency often refers to the faithful transmission of task-relevant forces to the user while maintaining stable interaction with the remote or virtual environment (*30, 31*). The current findings suggest the existence of an additional form of transparency: the device’s own mechanical presence must fade into the background of action, and that this form of transparency is not determined by physical mass alone. It also depends on whether the device’s forces are predictable and integrated into the user’s sensorimotor control system.

Perceived weight may therefore provide a useful behavioral metric for evaluating wearable haptic technologies. Traditional outcome measures such as task performance, comfort ratings, or embodiment questionnaires capture important aspects of use, but they may miss whether the device itself remains perceptually intrusive. A device can improve task performance while still feeling heavy, effortful, or foreign. Conversely, a device may become easier to use because its mass and contact forces are increasingly discounted. Measuring perceived device weight could therefore complement existing assessments of embodiment, usability, and haptic transparency (*13, 32, 33*).

These findings also connect to broader evidence that tools and artificial effectors can be integrated into bodily sensing. Tool-use studies show that external objects can extend somatosensory processing beyond the biological body (*24*), and sensorimotor predictions during tool use can attenuate self-generated tactile consequences (*28*). Supernumerary robotic limb studies similarly show that users can learn to coordinate artificial effectors with natural movement and that such use can alter neural body representation (*12, 34*). The present study adds that integration changes a basic perceptual property of the device itself: how heavy it feels.

From a design perspective, these results suggest that reducing physical mass is necessary but not sufficient to make a wearable device perceptually transparent. Designers should also consider how quickly the device comes to feel light during use. Predictable movement-to-feedback mappings, stable attachment, comfortable contact, and training protocols that expose users to the device’s altered dynamics may all promote sensorimotor transparency. This may be especially important for wearable haptic devices used in virtual reality, teleoperation, rehabilitation, or augmentation, where long-term acceptance depends on comfort and intuitive control as much as on technical performance (*2, 3*).

In conclusion, sensorimotor training reduced the perceived contribution of a wearable finger-extending exoskeleton to total effector weight. This effect was selective for the augmenting device rather than the biological hand, suggesting that perceived weight tracks the incorporation of artificial structures into sensorimotor control. For haptics and robotics, the central implication is that wearable devices should be evaluated not only by the sensations they deliver or the tasks they enable, but also by the extent to which their own mechanical presence becomes perceptually transparent.

## MATERIALS AND METHODS

### Participants

A total of 40 right-handed participants were tested. For the weight-perception analyses, data from 34 participants were included, with 17 participants in the exoskeleton group and 17 in the control group. The analyzed exoskeleton group had a mean age of 25.3 years (range = 18-61; 4 men, 13 women), and the analyzed control group had a mean age of 22.9 years (range = 18-43; 8 men, 9 women). Six participants were excluded before analysis because they consistently judged their hand as lighter than all presented comparison weights, preventing reliable estimation of psychometric functions and points of subjective equality. Importantly, these exclusions do not call into question the attenuation effect observed in the analyzed sample. Rather, these participants exhibited an extreme form of the same attenuation pattern, such that their perceived hand weight fell outside the measurable range of the task. Consequently, the reported attenuation likely provides a conservative estimate of the true magnitude of the effect.

All participants had normal or corrected-to-normal vision and reported no history of neurological or psychiatric conditions. Handedness was assessed by self-report and the Edinburgh Handedness Inventory. To ensure proper fitting of the exoskeletal device, participants were asked to trim their nails before testing. Recruitment took place through the Radboud University SONA system, and participants received course credits as compensation. The study was approved by the Ethics Committee of the Faculty of Social Sciences, Radboud University, and all participants provided written informed consent before participation.

### Devices

The wearable devices were custom-built passive finger exoskeletons mounted to the right hand. Each device consisted of four finger extensions and a thumb extension attached to an adjustable Velcro-secured wristband. In the augmentation condition, the device elongated each finger by approximately 10 cm while preserving native finger-joint mobility. The device was rigidly coupled to the distal phalanges by foam-padded fingertip caps, such that biological and artificial finger segments formed a single collinear mechanical module. Finger movements were transmitted directly to the extensions without additional joints or actuation.

The finger-extension device consisted of modular articulated segments terminating in a padded fingertip cap and external silicone gripping surface (Fig. 1B). Each artificial finger included a proximal segment with an elastic-band connection, an intermediate segment allowing adjustment to fingertip length, and a fingertip cap secured with a fastening screw and embedded nut. The internal surface of the cap contained foam padding attached with double-sided tape to provide a comfortable contact interface with the biological fingertip, whereas the outer silicone sleeve formed the contact surface used during object manipulation. This structure allowed the device to extend the functional reach of each finger while maintaining stable mechanical coupling to the participant’s hand during grasping and manipulation.

In the control condition, participants wore a structurally matched non-augmenting device with the same general attachment format, shape, fit, and interaction demands, but without meaningful finger extension. Devices were 3D-printed in PETG/PET and included a rigid internal core, a compliant silicone outer layer for object contact, foam-padded fingertip caps for comfort, and elastic tensioning elements to maintain stable coupling during movement. Two finger sizes, standard and large, were available to accommodate hand-size variability, along with multiple thumb widths to ensure proper fit. The standard and large exoskeletal finger components weighed 116 g and 134 g, respectively, with thumb components adding 28 g, 32 g, or 40 g depending on size. Control devices weighed 60 g or 67 g for the standard and large finger components, respectively, with thumb components adding 11 g, 13 g, or 15 g depending on size.

Each participant was individually fitted before testing by selecting the appropriate finger and thumb components, adjusting elastic tension to secure the finger caps, and verifying comfort and stability. During fitting, the fingertip caps were positioned so that the distal phalanges rested securely against the foam padding while still allowing participants to flex and extend their fingers comfortably. Participants reported any discomfort, pressure, or restriction of movement, and the device was adjusted when needed. After fitting, participants lifted a small ball using the device to confirm that it was securely attached and functionally effective before the experiment began.

### Experimental procedure

Participants completed four weight-perception measurements: hand only before training, hand plus device before training, hand plus device after training, and hand only after training. Between pre- and post-training testing, participants completed a 20- to 30-minute motor-training session while wearing their assigned device on the right hand. This order allowed assessment of perceived hand weight and combined hand-plus-device weight before and after sensorimotor experience with the device.

### Hand-weight measurement

Actual right-hand mass was estimated using a water-displacement method (*8*). Participants placed their right hand into a water-filled container positioned on a digital scale, submerging the hand up to the ulnar styloid process. The increase in scale reading was used as an estimate of the volume of water displaced by the hand. Because 1 g of water corresponds to approximately 1 cm^3^, the scale reading provided an estimate of hand volume. Three measurements were collected and averaged. Displaced water volume was converted to mass using a density coefficient of 1.09 g/cm^3^. For hand-plus-device conditions, the measured device mass was added to the estimated hand mass to obtain the physical mass of the effector.

### Weight-perception task

The procedure was adapted from an earlier study on self-estimation of hand weight (*8*). Participants were seated comfortably with their right hand hanging freely, while the left forearm rested on a surface at the same height with the palm facing downward. A wristband attached to the left wrist supported reference weights. Participants were blindfolded to eliminate visual cues. On each trial, a weight was suspended from the left wrist, and participants verbally judged whether it felt heavier or lighter than their right hand. In the hand-plus-device condition, judgments referred to the combined perceived weight of the hand and device.

At the start of each block, participants experienced the weight of the relevant right-hand effector. In the hand-only condition, this was the weight of the biological right hand. In the hand-plus-device condition, this was the combined weight of the right hand and attached device. Participants were instructed to use this experienced weight as the reference for subsequent comparison judgments. The right hand was then supported, and comparison weights were suspended from the wristband on the left wrist one at a time. The left wrist was used only to present comparison weights; participants’ judgments always referred to the weight of the right hand or right hand plus device.

Twenty logarithmically spaced reference weights ranging from 62 to 600 g were used. Stimulus selection followed an adaptive Bayesian staircase procedure (QUEST) (*35, 36*), with two interleaved staircases per block. One staircase was initialized below the expected perceived weight and the other above it, so that estimates could converge on the point of subjective equality from opposite directions. Inspection of the staircase trajectories confirmed that the upper- and lower-initialized staircases converged toward a common estimate, indicating that the final estimates were not driven by their initial starting values. On each trial, the QUEST algorithm selected the next comparison weight based on the participant’s previous responses. Participants responded verbally by saying whether the comparison weight felt lighter or heavier than the weight of the right hand or right hand plus device. No feedback was provided.

### Training procedure

Training was designed to familiarize participants with the device’s mechanical properties and support functional interaction during object manipulation. Participants performed grasping and manipulation tasks engaging different aspects of hand function, including peg transfer, color sorting, shape sorting, and Jenga-based tasks. Successful training was quantified using task-specific performance measures; pegboard transfer performance is presented in Fig. 3 as the primary training outcome, whereas the complete set of training-task outcomes is presented in Supplementary Fig. S1.

Performance was quantified using task-specific outcome measures, primarily completion time or, where applicable, the number of successfully completed actions within a fixed interval. Participants were instructed to use primarily the right, device-wearing hand, while the left hand was limited to minimal support, such as stabilizing objects. Tasks were repeated across two blocks to provide sustained practice. The same task set was used for both groups; the only difference was whether participants trained with the finger-extending exoskeleton or with the non-augmenting control device.

In the pegboard transfer task, participants transferred 14 PVC pegs between two boards using only the right hand fitted with the device, while the left hand was allowed only for stabilization. The task included large (9 cm) and small (5 cm) pegs, and the outcome measure was completion time. In the color-sorting task, participants sorted LEGO DUPLO bricks into trays according to color using the right hand; the outcome measure was completion time. In the shape-sorting task, participants inserted wooden shapes into corresponding slots in a shape-sorting box using the right hand, with the left hand permitted only for stabilization; the outcome measure was completion time. In the Jenga three-minute task, participants built as many complete floors of a Jenga tower as possible within three minutes using only the right hand; the outcome measure was the number of completed floors. In the full Jenga task, participants built an 18-floor Jenga tower using only the right hand; the outcome measure was total completion time.

### Psychometric estimation

For model-based analyses, attenuation parameters were instead estimated via refitting of psychometric functions to trial-level data, using the attenuation model described in the Results (see *Theoretical model of weight attenuation*). In this approach, perceived weight was modeled directly as a function of actual effector mass, and no additional normalization step was applied.

For ease of interpretation and consistency with condition-level analyses, attenuation parameters estimated from the model (α) were converted to percentage attenuation values by multiplying parameter values by 100. In this framework, positive values indicate underestimation of weight and negative values indicate overestimation. Thus, all reported attenuation values reflect percentage deviation from veridical perception.

### Regression-based decomposition of attenuation parameters

The primary analysis in the main text used a regression-based framework applied to participant-level attenuation parameters. This framework decomposed perceived weight attenuation into three components: (1) baseline attenuation, (2) a general training effect, and (3) a device-specific training effect. Perceived weight was modeled as an attenuated function of physical mass, parameterized by attenuation coefficients. For reporting purposes, all regression coefficients corresponding to attenuation parameters were expressed in percentage attenuation units by multiplying the original parameter values by 100.

The attenuation parameters were decomposed into their component terms using the following two-step procedure applied to each participant’s judgments.

#### Step 1: Extract the attenuation parameters from participant judgments

We used model-driven curve fitting to estimate all attenuation parameters (Equations 1-2) from each participant’s data. We used Cumulative Normal functions to refit their judgments made during the staircase procedure. These functions modelled the probability that the participant reported that the weight of the comparison object was greater than their hand or hand+exoskeleton weight,

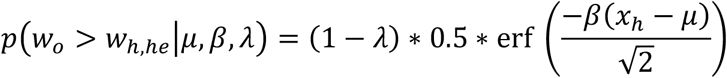

where *w*_*o*_ is the weight of the comparison object, erf is the complementary error function, *λ* is the lapse rate, *β* is the slope, and *μ* is the point of subjective equality (PSE). In the case of our experiment, the PSE is equivalent to the perceived weight of the hand. Therefore, by substituting *μ* with Equations 1 and 2 (i.e., 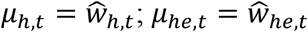), we can fit the hand and hand+exoskeleton blocks to extract *α*_*h,t*_ and *α*_*e,t*_.

Fitting the entire dataset simultaneously involved nine free parameters with the following constraints: all four attenuation parameters *α*_*h,t*_ and *α*_*e,t*_ [-1, 1]; four block-specific slopes, *β* [0, 1], and a single lapse rate *λ* [0, 0.1]. All curve-fitting was performed with the Palamedes Toolbox in Matlab 2019b as well as custom scripts for constrained parameter optimization (using *fmincon*).

#### Step 2: Group-level regression to decompose attenuation parameters

We used group-level regression to decompose the attenuation parameters in Step 1 into all variables from Equation 3. This was done separately for *α*_*h*_ and *α*_*e*_ by maximizing the likelihood of the attenuation data given the parameters of the following regression equation.

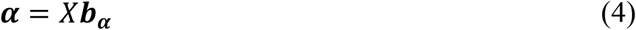

Where ***α*** is a vector of 68 attenuation parameters (17 participants per two conditions, each with two time points); ***b***_***α***_ is a vector of the attenuation variables (Equation 3), ***b***_***α***_ = [*α*_1_, *δ*_*g*_, *δ*_*et*_]^*T*^; and *X* is a matrix of dummy variables that enable the decomposition—*X*[, 1] is a vector of ones for all data points; *X*[, 2] is a vector whose values equal zero for all pre-training datapoints and equal one for all post-training datapoints; and, *X*[, 3] is a vector whose values equal one for all post-exoskeleton training datapoints and zero for all others. To best account for individual-participant variability, the regressions were fit using linear mixed effects modelling with ***b***_***α***_ as fixed effects, participant as a random intercept, and time (pre-post) as a random slope. Fixed-effect tests were obtained from the corresponding linear mixed-effects models, with degrees of freedom estimated using the residual degrees-of-freedom method.

### Bayesian statistical analyses

We first used a Bayesian bootstrapping procedure to estimate posterior distributions of each attenuation variable. Distributions were created from 5000 resamples. Per resample, we weighted each attenuation estimate’s contribution to the model fit using values drawn from a Dirichlet distribution. This was done simultaneously for the hand and exoskeleton parameters, ensuring that we could also derive posterior distributions of the differences between hand and exoskeleton parameter values.

Kernel density estimation was applied to the histogram of resamples to derive a probability density function for the posterior distribution, *P*_*b****α***_. This procedure thus resulted in 9 posterior distributions, six distributions of attenuation variables (3 per hand and exoskeleton) and three distributions of the differences between hand and exoskeleton.

We then statistically assessed whether each distribution differed significantly from zero. To do so, we calculated Bayes factors for each variable and their difference distributions: 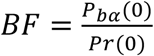, where *P*_*b****α***_(0) is the probability of zero given the posterior distribution and *Pr*(0) is the prior probability of zero given a Cauchy distribution centered on zero. To determine the robustness of our results, we derived Bayes factors for several prior widths ranging from 0.2-1.0.

## Acknowledgments

We thank Sibrecht Bouwstra for her key role in the development of the exoskeletons and the Technical Support Group of the Donders Centre for Cognition for additional support.

## Author contributions

Conceptualization: DR, AS, MRL, LEM

Methodology: DR, AS, MRL, LEM

Investigation: DR, AS, LEM

Visualization: DR, AS, LEM

Supervision: DR, LEM

Writing - original draft: DR

Writing - review & editing: DR, LEM, MRL

## Data, code and materials availability

The data and code that support the findings of this study will be available from the Open Science Framework upon publication. No new materials were generated in this study.

