## Supplementary Materials for "Sensorimotor training lightens the perceived weight of body augmentation devices"

Dominika Radziun* *et al.*

**This PDF file includes:**

Supplementary Results

### Supplementary Results

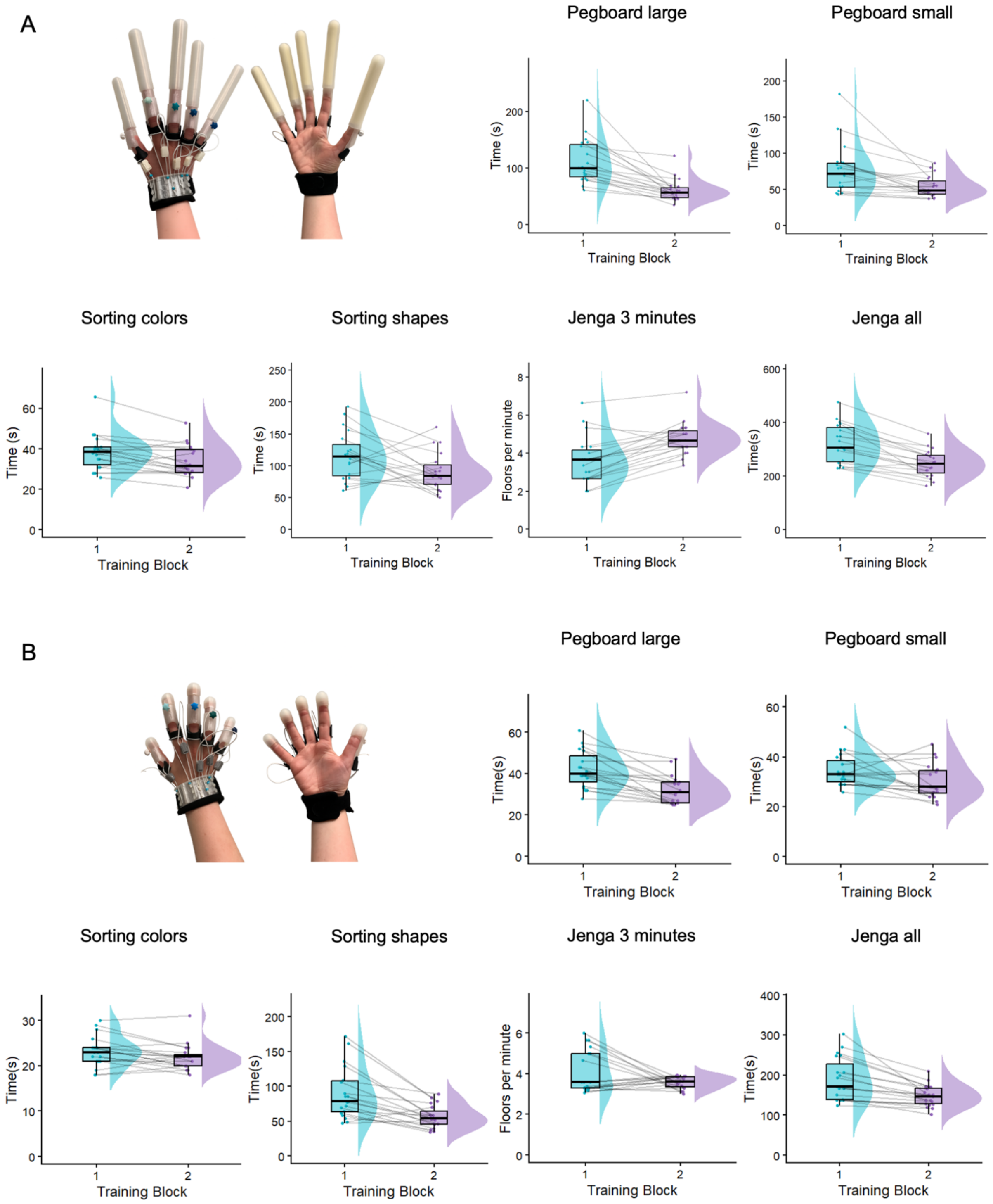


**Figure S1.** Overview of training tasks. In A, participants trained with the finger-extending exoskeleton; in B, participants trained with a control device. Each panel displays two rows of task conditions: the top row shows the pegboard transfer tasks with large and small pegs, and the bottom row shows the sorting colors task, sorting shapes task, the Jenga 3-minute task, and the Jenga all task.

**Performance on additional training tasks**

Participants completed two blocks of object-manipulation training while wearing either the finger-extending exoskeleton or the structurally matched control device. In addition to the pegboard-transfer tasks reported in the main text, training included color sorting, shape sorting, a three-minute Jenga task, and construction of a complete Jenga tower.

In the exoskeleton group, completion times decreased from the first to the second training block for color sorting (Wilcoxon $V = 23.5$, $p = 0.012$, $BF_{10} = 8.16$), shape sorting ($t(19) = 2.56$, $p = 0.019$, $BF_{10} = 3.00$), and full Jenga construction ($t(19) = 7.35$, $p < 0.001$, $BF_{10} = 30028.86$). Participants also completed more Jenga floors during the three-minute task in the second block than in the first ($t(19) = 5.71$, $p < 0.001$, $BF_{10} = 1405.76$).

The control group likewise showed reduced completion times for color sorting (Wilcoxon $V = 37.0$, $p = 0.019$, $BF_{10} = 4.02$), shape sorting ($t(19) = 4.82$, $p < 0.001$, $BF_{10} = 239.71$), and full Jenga construction ($V = 0.0$, $p < 0.001$, $BF_{10} = 584.72$). However, the number of floors completed during the three-minute Jenga task did not reliably change across training blocks ($V = 89.0$, $p = 0.571$, $BF_{10} = 0.81$). Together, these results indicate that participants generally improved their control of both devices during active object manipulation, with particularly strong evidence of improvement across the Jenga tasks in the exoskeleton group.
